# Antibiotic-Induced Gut Dysbiosis Differentially Alters Hippocampal Glial Proliferation and Response After TMEV Infection

**DOI:** 10.1101/2025.05.23.655804

**Authors:** Sophia Shonka, Inga Erickson, Melissa Barker-Haliski

**Affiliations:** Department of Pharmaceutics, University of Washington, Seattle, WA

**Keywords:** Carbamazepine, neuroinflammation, neurodegeneration, microbiome, astrogliosis, microgliosis

## Abstract

Viral encephalitis is a major worldwide cause of acquired epilepsy, yet the impact of environmental and non-neuronal factors on an individual’s risk for epilepsy is understudied. For example, the gut microbiome influences immune system function yet the impact of the gut microbiome on seizure-associated neuropathology remains poorly understood. Using the Theiler’s murine encephalomyelitis virus (TMEV) mouse model of infection-induced acute symptomatic seizures (ASyS), we aimed to investigate how antibiotic (ABX)-induced gut dysbiosis during a brain viral infection could influence resulting hippocampal neuropathology. Further, we included co-administration of carbamazepine (CBZ) to assess the extent to which antiseizure medicines could also shift the neurological impact of ASyS and gut dysbiosis. Brain tissue from TMEV-infected mice with and without gut dysbiosis was assessed for neurodegeneration and glial response (astrocytes and microglia). TMEV infection primarily drove neuroinflammatory changes in CA1, including increased astrogliosis, microgliosis, and microglial activation. ABX-induced dysbiosis exacerbated neuroinflammation across hippocampal subregions, markedly increasing microgliosis in CA3 and DG, and elevating microglial and neuronal proliferation in CA1. TMEV infection-induced astrogliosis was impacted by dysbiosis in a region-dependent manner, being worsened in DG while being alleviated in CA3. CBZ was neuroprotective selectively within DG, reducing neurodegeneration and microglial immunoreactivity with dysbiosis. Astroglial proliferation occurred regardless of gut microbiome integrity. Altogether, gut dysbiosis shapes hippocampal neuroimmune responses following viral infection-induced ASyS in a region-dependent manner, and CBZ may confer a neuroprotective effect. Together this work highlights the acute neuroinflammatory impact of infection-induced ASyS and reveals an underappreciated contribution of the gut-brain-axis to seizure-related neuropathology.

## Introduction

Epilepsy is an underrecognized long-term complication of central nervous system (CNS) infection-induced encephalitis (Singh et al., 2008), increasing epileptogenesis risk up to 22-fold (Libbey and Fujinami, 2011; Misra et al., 2008). Importantly, low sociodemographic index nations carry a higher burden of CNS infection-induced acquired epilepsy (e.g., 37% of epilepsy cases in Mali can be attributed to brain infection (Ba-Diop et al., 2014) vs 5.8% in the United States (Bosak et al., 2019). Thus, viral infection-induced encephalitis is a unique risk factor that disproportionately contributes to the global burden of epilepsy. However, rodent models of encephalitis-induced acute symptomatic seizures (ASyS) and acquired epilepsy are less frequently integrated into preclinical epilepsy research. One translationally relevant rodent model of infection-induced acute seizures and acquired epilepsy is induced through Theiler’s murine encephalomyelitis virus (TMEV) infection directly into the CNS. Hippocampal TMEV infection leads to frequent ASyS in C57BL/6J mice in a viral titer-dependent manner, with up to 50% of mice with acute seizure presentation going on to later develop spontaneous recurrent seizures (SRS) (DePaula-Silva et al., 2017; Libbey and Fujinami, 2011; Stewart et al., 2010a). Mice also display similar behavioral comorbidities (Barker-Haliski et al., 2022, 2016, 2015; Umpierre et al., 2014), and neuropathology (Barker-Haliski et al., 2015; Libbey et al., 2016; Libbey and Fujinami, 2011; Loewen et al., 2016; Umpierre et al., 2014) observed in clinical cases of infection-induced acquired temporal lobe epilepsy (TLE). Further, the TMEV model is especially advantageous due to the lower mortality rate than that which is observed in other viral encephalitis-induced epilepsy models (DePaula-Silva et al., 2017; Libbey and Fujinami, 2011). Thus, the TMEV mouse model is an etiologically relevant model of viral infection-induced ASyS and chronic SRS that reproduces multiple clinical features of acquired TLE. For this reason, the TMEV model of ASyS may fill an important translational research gap to address the global epilepsy burden.

Neuroinflammation is a key mediator of acute and chronic seizures (Gaspard et al., 2018; Goyal et al., 2025; Guillemaud et al., 2025; Misra et al., 2008; Sakuma et al., 2015) and neuropathology following viral encephalitis (Loewen et al., 2016; Umpierre et al., 2014). However, emerging evidence suggests that peripheral immune regulators, particularly the gut microbiome (Agirman et al., 2021; Ding et al., 2021; L. Liu et al., 2022), play an important, yet understudied, role in shaping these neuroimmune responses. Clinically, changes in gut microbiome composition elevate seizure risk (Kwack et al., 2024) and drug-resistant epilepsy patients also exhibit compositional differences in the intestinal microbiome when compared to drug-sensitive patients (Cheraghmakani et al., 2021; Peng et al., 2018), highlighting a bidirectional cross-talk between the gastrointestinal tract and the brain. Moreover, we previously demonstrated that experimentally-induced intestinal dysbiosis in the TMEV model is sufficient to alter ASyS burden and dramatically alter CBZ antiseizure activity (Erickson et al., 2025). In our earlier study, CBZ was anticonvulsant under eubiotic conditions, with CBZ-treated mice experiencing significantly fewer seizures during the acute infection period compared to vehicle-treated mice (Supplementary Figure 1). However, following antibiotic (ABX)-induced dysbiosis, CBZ displayed proconvulsant effects with significantly more seizures than vehicle-treated mice with evoked gut dysbiosis. Notably, plasma concentration of CBZ did not differ between dysbiotic and eubiotic mice (Supplementary Figure 2), suggesting that experimentally evoked gut dysbiosis did not impact the pharmacokinetics of CBZ (Erickson et al., 2025). Together, these findings position the gut microbiome as a key determinant of neuroimmune signaling, which may carry downstream effects on antiseizure medication (ASM)-mediated seizure control in the setting of viral infection-induced ASyS presentation. However, few studies have fully explored the bidirectional interactions between the gut-brain axis and ASM treatment on seizure-induced hippocampal neuroinflammation and neuropathology. Therefore, we hypothesized that our previously observed effects of gut dysbiosis on ASyS burden and CBZ efficacy (Erickson et al., 2025), could alter TMEV infection-induced neuroinflammation, including astrogliosis and microgliosis.

The gut microbiome is an understudied contributor to seizures in epilepsy and therapeutic response, and how experimentally evoked gut dysbiosis influences neuropathological changes with TMEV infection-induced ASyS is unclear. We thus sought to extend our earlier study to assess the effects of gut dysbiosis, ASyS history, and CBZ administration (Erickson et al., 2025) on neuroinflammatory hallmarks associated with CNS infection and viral encephalitis-induced seizures. We herein further demonstrate that the gut microbiome shifts ASyS-induced neuropathological damage after brain infection with TMEV in mice. Notably, we show that gut dysbiosis differentially influences gliosis across hippocampal regions after viral infection and ASyS presentation. Altogether, this study adds to mounting evidence that the gut microbiome plays an outsized role in seizure-related neuroinflammatory impacts within the rodent hippocampus, which may drive susceptibility to future chronic SRS onset and epileptogenesis.

## Experimental Procedures

### Animals

Mouse brain tissue from our previously published study was used, with detailed experimental methods described in that earlier publication (Erickson et al., 2025) and summarized in Figure 1. Briefly, male C56BL/6J mice (4-5 weeks old; Jackson Labs) were pretreated with an oral antibiotic cocktail (ABX; ampicillin (1 g/L), metronidazole (1 g/L), neomycin sulfate (1 g/L), and vancomycin (0.5 g/L) dissolved in saline) or saline (SAL) once daily for 10 days beginning upon arrival until conclusion of in-life testing 7 days post infection (Figure 1). Two days following arrival, mice were either infected with TMEV (20 μL, 3.0×10^5^ PFU, intracerebrally) or underwent sham infection with sterile PBS. TMEV titers were isolated from stock originally provided by Robert Fujinami at the University of Utah. Then, during the acute infection period (post-infection days 3-7), CBZ (20 mg/kg) or vehicle (VEH; 0.5% methylcellulose) was administered intraperitoneally twice daily 30 minutes prior to each twice-daily handling-induced seizure assessment (minimum 4 hours between sessions). Twice-daily seizure assessment involved brief cage agitation (<30 seconds), followed by individual handling of mice to visually assess for evoked seizure onset. Typically, evoked secondarily generalized seizures occur in susceptible animals within this 5–10-minute observation period. The presence and severity of handling-induced seizures was scored according to the Racine scale (1 – mouth and facial clonus; 2 – head bobbing; 3 – single forelimb clonus; 4 – bilateral forelimb clonus plus rearing; 5 – stage 4 plus repeated rearing and falling), as previously reported (Libbey and Fujinami, 2011). On the 7^th^ day after TMEV infection, all mice were euthanized and tissues collected for immunofluorescence processing as previously detailed (Erickson et al., 2025). All animal use was approved by the University of Washington Institutional Animal Care and Use Committee (protocol 4387-02) and conformed to ARRIVE guidelines (Sert et al., 2020).

**Figure 1.**
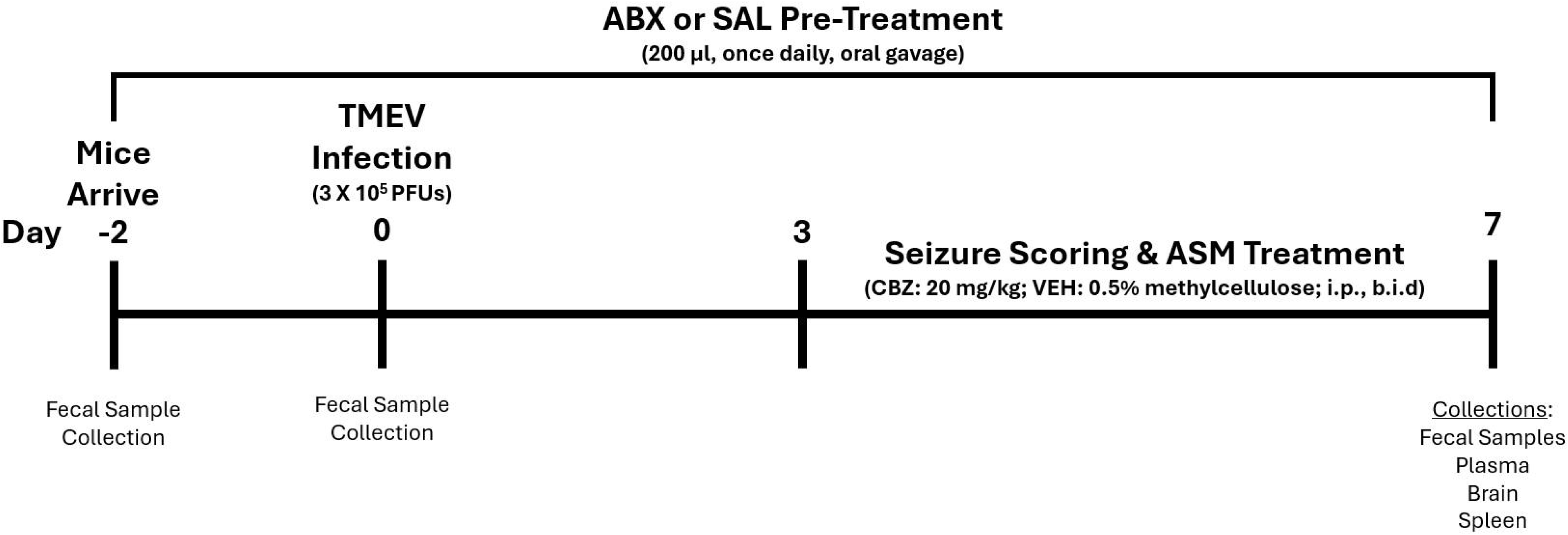
Overview of study timeline to test the hypothesis that experimentally induced alterations in the gut microbiome using antibiotics (ABX) would evoke significant differences in acute behavioral seizure presentation, anticonvulsant activity, and neuroinflammatory changes with the Theiler’s murine encephalomyelitis virus (TMEV) model of acute infection-induced acute seizures in male C57BL/6J mice (aged 4-5 weeks). The antibiotic cocktail (ABX) consisted of ampicillin (1 g/L), metronidazole (1 g/L), neomycin sulfate (1 g/L), and vancomycin (0.5 g/L) administered 1x/day by oral gavage; half of mice were randomized to saline pretreatment. The dose of carbamazepine (CBZ) was selected based on its known anticonvulsant activity in a variety of rodent seizure models. CBZ (20 mg/kg, i.p.) was administered to mice twice per day 30 min prior to behavioral seizure scoring. Mice were randomized to receive either 0.5% methylcellulose vehicle (VEH) or CBZ from day 3 post TMEV infection. On the final testing day, animals were euthanized at discrete time points after CBZ or VEH administration (0, 15, or 60 min post-CBZ administration) for a limited pharmacokinetic study of CBZ plasma concentrations and to assess potential that ABX-induced gut microbiome dysbiosis could influence neuroinflammation 7 days after TMEV infection.

### Immunofluorescence

Brain tissue was cryosectioned (Leica) at a thickness of 40 µm, mounted on Fisher Superfrost Plus microscope slides (1255015; 4 sections/slide), and stored at 4 C. Astroglial and neuronal proliferation in the hippocampus (CA1, CA3, and dentate gyrus [DG]) was assessed using immunofluorescence (IF) procedures. Slides were washed three times with 0.1 M PBS and blocked in a Triton X solution (0.1% Triton and 1% bovine serum albumin [BSA] in 0.1 M PBS) with 10% goat serum for 2 hours. Tissue was incubated overnight at 4 C with primary antibody directed to Ki-67 (Abcam ab16667; 1:300) diluted in the same Triton X solution used for blocking step. Sections were then rinsed three times in 0.1 M PBS and incubated for 1 hour in goat anti-rabbit secondary conjugated to Alexa Fluor 555 (Abcam ab150078; 1:500) with anti-GFAP conjugated to Alexa Fluor 488 (Sigma MAB3402X; 1:1000) or anti-NeuN conjugated to Alexa Fluor 488 (Sigma MAB377X; 1:300). Slides were then fixed with Prolong Gold with DAPI nuclear counterstain (Invitrogen P36935).

Finally, microglia proliferation and activation in the hippocampus were assessed using similar IF procedures as for astroglia. Slides were washed three times with 0.3% Triton X in 0.1 M PBS and blocked for 1 hour with the same Triton X solution and 5% BSA. Tissue was incubated overnight at 4 C with primary antibodies directed to Ki-67 (ThermoFisher 14-5698-80; 1:300), Iba-1 (Fujifilm 019-19741; 1:1000), and CD68 (ThermoFisher 968-MSM3-P0; 1:1000) diluted with 1% BSA and the 0.3% Triton X solution. Slides were then rinsed in the 0.3% Triton X solution three times and incubated for 2 hours in, respectively, goat anti-rat secondary conjugated to Alexa Fluor 488 (ThermoFisher A-11006; 1:500), goat anti-rabbit conjugated to Alexa Fluor 555 (Abcam ab150078; 1:1000), and goat anti-mouse conjugated to Alexa Fluor 647 (ThermoFisher A-21235; 1:2000). Slides were then fixed with Prolong Gold with DAPI nuclear counterstain (Invitrogen P36935). Sample sizes varied across experimental groups and stains due to occasional tissue loss during histological processing (Table 1).

**Table 1.** Sample size summary and ImageJ analysis parameters for histological analysis in hippocampus of male C57BL/6J mice with and without experimentally evoked intestinal dysbiosis (SAL or ABX) 7 days after an acute TMEV or sham infection and subsequent administration of the antiseizure medicine, carbamazepine (CBZ, 20 mg/kg, i.p., bid). Due to tissue loss during histological staining, sample sizes were inconsistent across groups and stains. Only mice that displayed at least one 3+ Racine seizure following TMEV infection were included in data analysis, except for the Iba-1 and CD68 analysis where the experimental group sample size for dysbiotic mice did not include enough mice that displayed severe seizures so all dysbiotic mice were included in this analysis regardless of seizure severity. The threshold method utilized in ImageJ for stain signal quantification is also indicated below, with Li being the name of one threshold method.

|  | FJC | Iba-1 & Ki-67 | GFAP &<br>Ki-67 | NeuN &<br>Ki-67 | Iba-1 & CD68 |
| --- | --- | --- | --- | --- | --- |
| Experimental Group N (3+ Racine seizure N, % with 3+ Racine seizure) |  |  |  |  |  |
| Sham-SAL-VEH | 13 (0, 0%) | 12 (0, 0%) | 11 (0, 0%) | 12 (0, 0%) | 10 (0, 0%) |
| Sham-ABX-VEH | 1 (0, 0%) | 3 (0, 0%) | 2 (0, 0%) | 2 (0, 0%) | 0 (0, 0%) |
| TMEV-SAL-VEH | 9 (5, 56%) | 14 (10, 71%) | 13 (9, 69%) | 14 (9, 64%) | 9 (5, 56%) |
| TMEV-SAL-CBZ | 11 (3, 27%) | 12 (0, 0%) | 9 (2, 22%) | 10 (2, 20%) | 6 (0, 0%) |
| TMEV-ABX-VEH | 12 (2, 17%) | 14 (3, 21%) | 12 (2, 17%) | 13 (3, 23%) | 9 (0, 0%) |
| TMEV-ABX-CBZ | 9 (4, 44%) | 10 (5, 50%) | 10 (6, 60%) | 8 (5, 63%) | 6 (3, 50%) |
| Threshold Method | Moments | Iba: Li<br>Ki67: Mean | Percentile | Percentile | Iba: Li<br>CD68: Percentile |

### Microscopy and Image Analysis

All slides were imaged on an upright fluorescent microscope (Leica DM6 B) with a 20x objective with constant acquisition settings (100 ms exposure, 1x gain). Raw grey scale 16-bit images from the Iba-1, GFAP, FJC, and NeuN channels were utilized to measure the percent field area of stain-positive signal using ImageJ Fiji (1.54p) in order to identify the impact of TMEV, ABX-induced gut dysbiosis, and CBZ on microglia, astroglia, cell degeneration, and neurons, respectively. Then the percent field area of Iba-1-, GFAP-, and NeuN-positive signal co-labeled with Ki-67 was measured using an ImageJ colocalization plugin (https://imagej.net/ij/plugins/colocalization.html) to quantify the impact of TMEV, ABX-induced gut dysbiosis, and CBZ on microglial, astroglial, and neuronal proliferation, respectively. Finally, the percent field area of Iba-1-positive signal co-labeled with CD68 was also measured using the colocalization plugin to identify the impact of TMEV, ABX-induced gut dysbiosis, and repeated CBZ administration on microglial activation. All raw images were filtered using Gaussian Blur (sigma = 3), and threshold applied using stain-specific settings (Table 1), prior to the particle analysis measuring the percent field area of the stain signal or co-labeled stain signal (particle size range: 0-Infinity; Circularity range: 0.5-1). Stain-specific threshold methods and ranges were determined using a subset of the images used for analysis were selected at random for each stain (12 images total, 6 images/infection group, 4 images/HPC region). The auto threshold function was utilized for all subset images to determine the default threshold range for each threshold method. The threshold method with the lowest threshold range standard deviation was utilized for image analysis using the default threshold range for each image.

### Statistical Analysis

Two-way ANOVAs with Tukey *post-hoc* t-tests were used to determine the impact of TMEV infection (TMEV vs Sham) and dysbiosis status (SAL vs ABX) on percent field area of FJC-, Iba-1, GFAP-, and NeuN-positive signal as well as the percent co-labeled field area of Iba-1, GFAP, and NeuN with Ki-67. Due to limited sample sizes, the percent co-labeled field area of Iba-1 with CD68 was analyzed using t-tests to determine the impact of TMEV infection (TMEV vs Sham) in eubiotic mice. Next, two-way ANOVAs with Tukey *post-hoc* t-tests were used to determine the impact of dysbiosis (SAL vs ABX) and ASM treatment (VEH vs CBZ) in TMEV mice on percent field area of GFAP- and NeuN-positive signal, as well as the percent co-labeled field area of GFAP and NeuN with Ki-67. Due to limited sample sizes, the percent field area of Iba-1, and percent co-labeled field area of Iba-1 with Ki67 and Iba-1 with CD68 was analyzed using t-tests to determine the impact of ASM treatment (VEH vs CBZ) in dysbiotic mice with TMEV. Only mice that displayed a 3+ Racine seizure during the acute infection period were included in analysis except for the t-test analyzing the impact of ASM treatment on the percent co-labeled field area of Iba-1 with CD68, whereby sample size limitations prevented the exclusion of mice without 3+ Racine seizures. Hippocampal regions (CA1, CA3, and DG) were analyzed separately for each analysis. Statistical analysis was performed in GraphPad Prism v11 or later, with p < 0.05 significant.

## Results

Considering the measurable microgliosis that occurs with TMEV infection and ASyS presentation (Loewen et al., 2016), we evaluated microglial phenotypes in hippocampal subregions in dysbiotic and eubiotic mice with and without TMEV infection, including microglia, microglial proliferation, and microglial activation. TMEV selectively increased total field area with Iba-1 positive signal area in CA1 (F(1,24)=6.12, p<0.05; Figure 2A), while ABX-induced gut dysbiosis increased microglia area in CA3 (F(1,24)=14.28, p<0.001; Figure 2B) and DG (F(1,24)=18.57, p<0.001; Figure 2C). However, CBZ only reduced microglia area in dysbiotic mice in DG (t(5.71)=4.21, p<0.01; Figure 2F). *Post hoc* comparisons revealed greater microglia immunoreactivity in dysbiotic TMEV mice versus eubiotic sham mice across hippocampal regions (CA1: p<0.05, CA3: p<0.01, DG: p<0.01). Further, dysbiotic sham mice showed greater microgliosis in DG versus eubiotic sham mice (p<0.01), and eubiotic TMEV mice showed greater microgliosis in CA1 compared to eubiotic sham mice (p<0.01). There was also a trending increase in microglia area in dysbiotic versus eubiotic TMEV-infected mice (p=0.05). Thus, TMEV and ABX-induced dysbiosis differentially increased total field area with Iba-1 positive signal across hippocampal regions while CBZ administration was associated with selective protective effects against ABX-induced increases in microglia area in TMEV mice. Meanwhile, ABX-induced dysbiosis increased microglial proliferation in CA1 (F(1,24)=6.11, p<0.05; Figure 3A) but CBZ did not protect against the observed elevation in microglial proliferation (Figure 3D). No other significant effects were observed on microglial proliferation. Like the microgliosis results, microglial activation selectively increased in TMEV mice in CA1 (t(6.48)=2.98, p<0.05; Figure 4A). However, no other significant results were found and CBZ made no significant impact on microglial activation. Together these results demonstrate that TMEV primarily affects microglia in CA1, increasing both microgliosis and microglial activation, while ABX-induced dysbiosis elevates microgliosis in CA3 and DG, and microglial proliferation in CA1. Thus, our study demonstrates hippocampal region-dependent effects of TMEV and ABX-induced dysbiosis on microglia activation and response.

**Figure 2.**
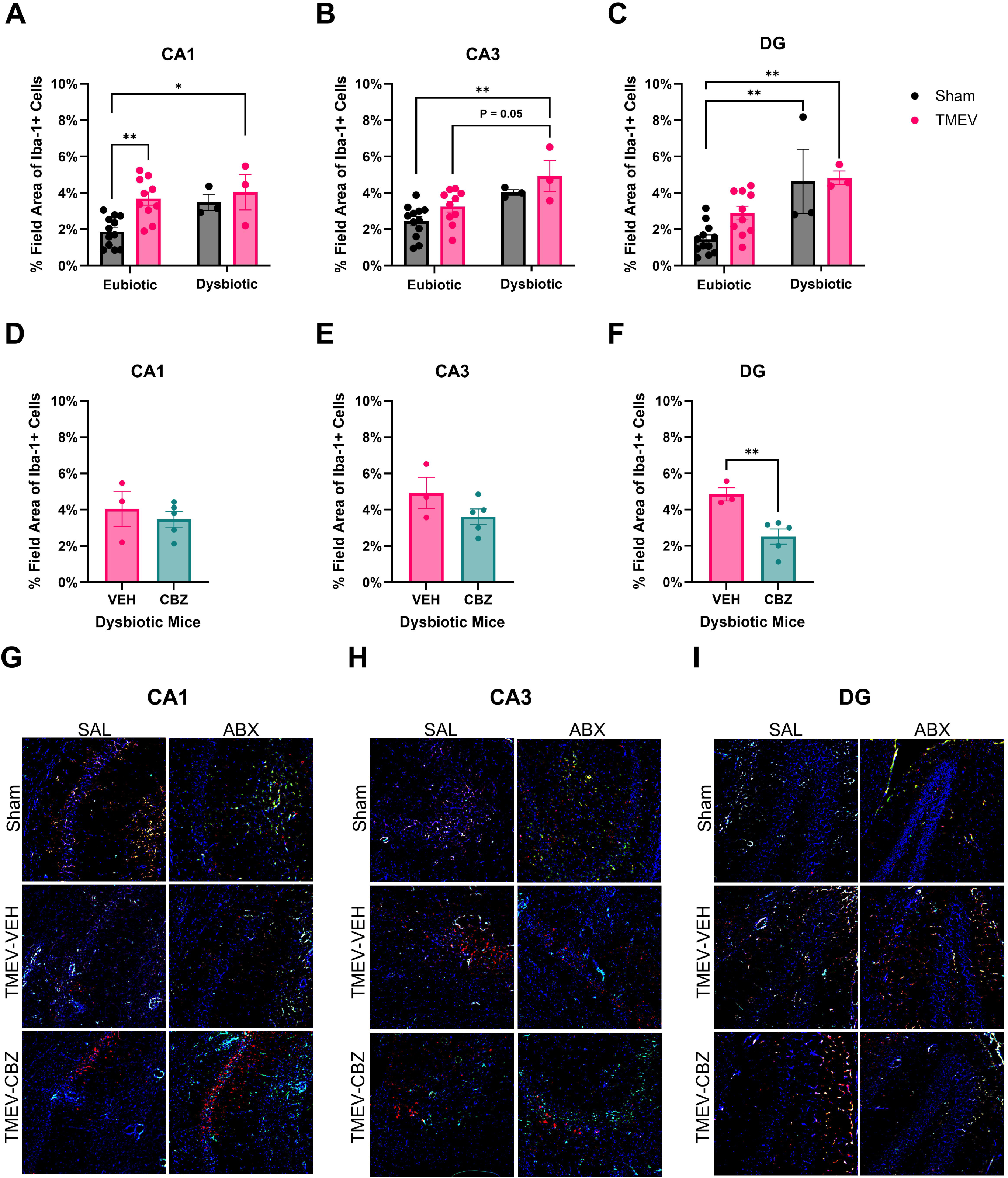
Brain infection with the neurovirulent pathogen TMEV leads to significant changes in the total field area of dorsal hippocampus of male C57BL/6J mice with microglial marker-positive total field area. C57BL/6J mice with and without experimentally evoked intestinal dysbiosis (SAL or ABX) were assessed 7 days after an acute TMEV or sham infection and subsequent administration of the antiseizure medicine, carbamazepine (CBZ, 20 mg/kg, i.p., bid). Only mice that experienced at least one 3+ Racine seizure following TMEV infection were included in data analysis. A) TMEV infection significantly increased microglial percent field area in the CA1 hippocampal subregion 7 days post-infection. B) Experimentally evoked gut dysbiosis increased percent field area with microglia-specific antibody marker signal (Iba-1), specifically in TMEV-infected mice, in the CA3 hippocampal subregion 7 days post-infection. C) Experimentally evoked gut dysbiosis increased percent field area with microglia-specific antibody marker signal (Iba-1), regardless of infection status, in the DG hippocampal subregion 7 days post-infection. D-E) CBZ treatment had no significant impact on microglial-specific percent field area 7 days post-infection in mice with experimentally-evoked dysbiosis. G-I) Representative photomicrographs of the CA1 (G), CA3 (H), and DG (I) hippocampal subregions of Iba-1+ (red), Ki-67+ (green), and CD68+ (teal) cells with DAPI nuclear counter stain (blue). *, *p* < 0.05, **, *p* < 0.01. Graphs show mean ± SEM, scale bar = 100 µm.

**Figure 3.**
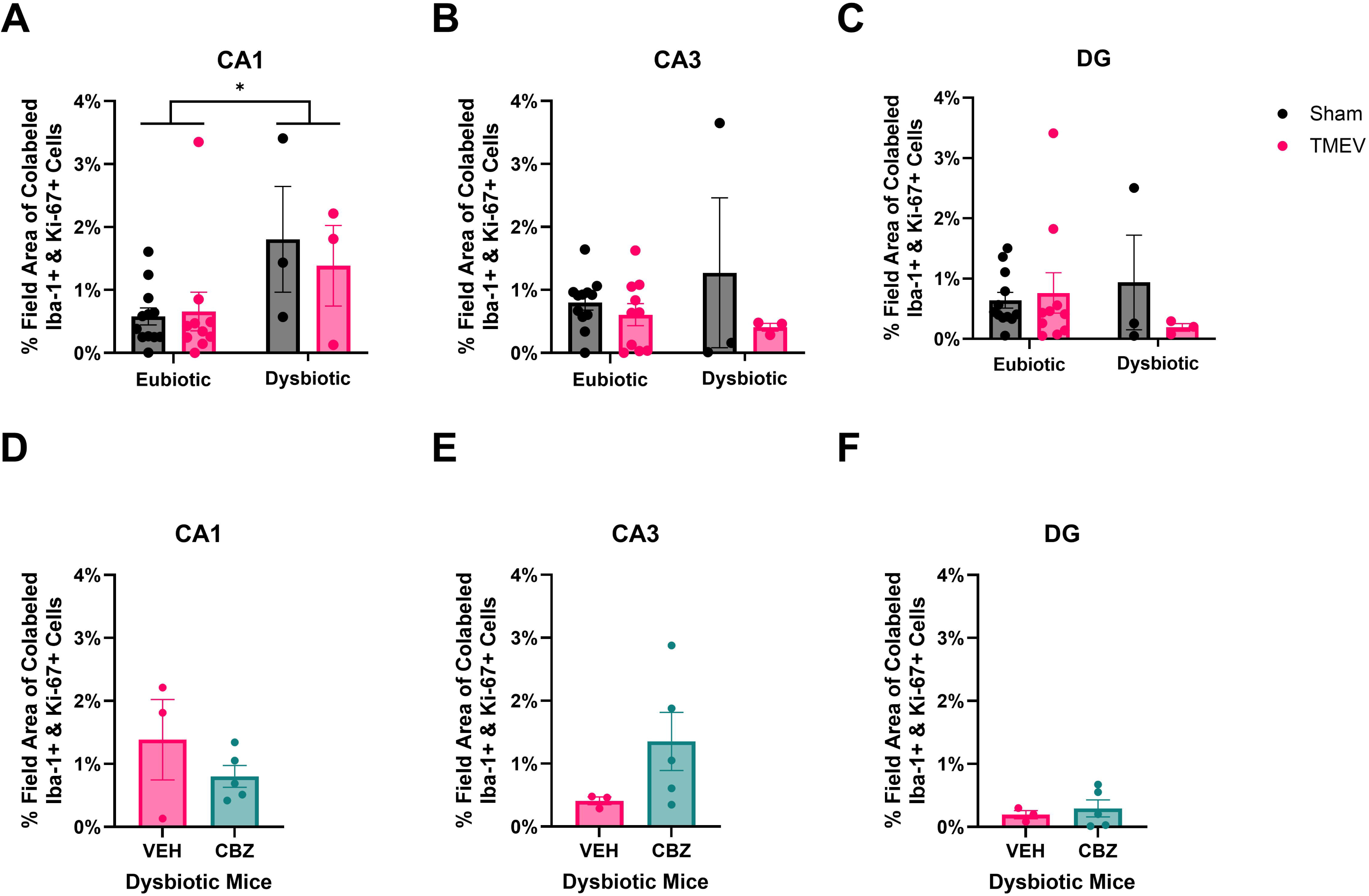
Microglial proliferation field area changes in hippocampus of male C57BL/6J mice with and without experimentally evoked intestinal dysbiosis (SAL or ABX) 7 days after an acute TMEV or sham infection and subsequent administration of the antiseizure medicine, carbamazepine (CBZ, 20 mg/kg, i.p., bid). Only mice that experienced at least one 3+ Racine seizure following TMEV infection were included in data analysis. A) Experimentally evoked gut dysbiosis increased microglial proliferation regardless of infection status 7 days post-infection in the CA1 hippocampal subregion. B-C) Neither TMEV infection nor experimentally evoked gut dysbiosis impacted microglial proliferation in the CA3 (B) or DG (C) hippocampal subregions 7 days post-infection. D-F) CBZ treatment did not impact microglial proliferation in the CA1 (D), CA3 (E), or DG (F) hippocampal subregions 7 days post-TMEV infection in mice with experimentally evoked gut dysbiosis. *, *p* < 0.05. Graphs show mean ± SEM.

**Figure 4.**
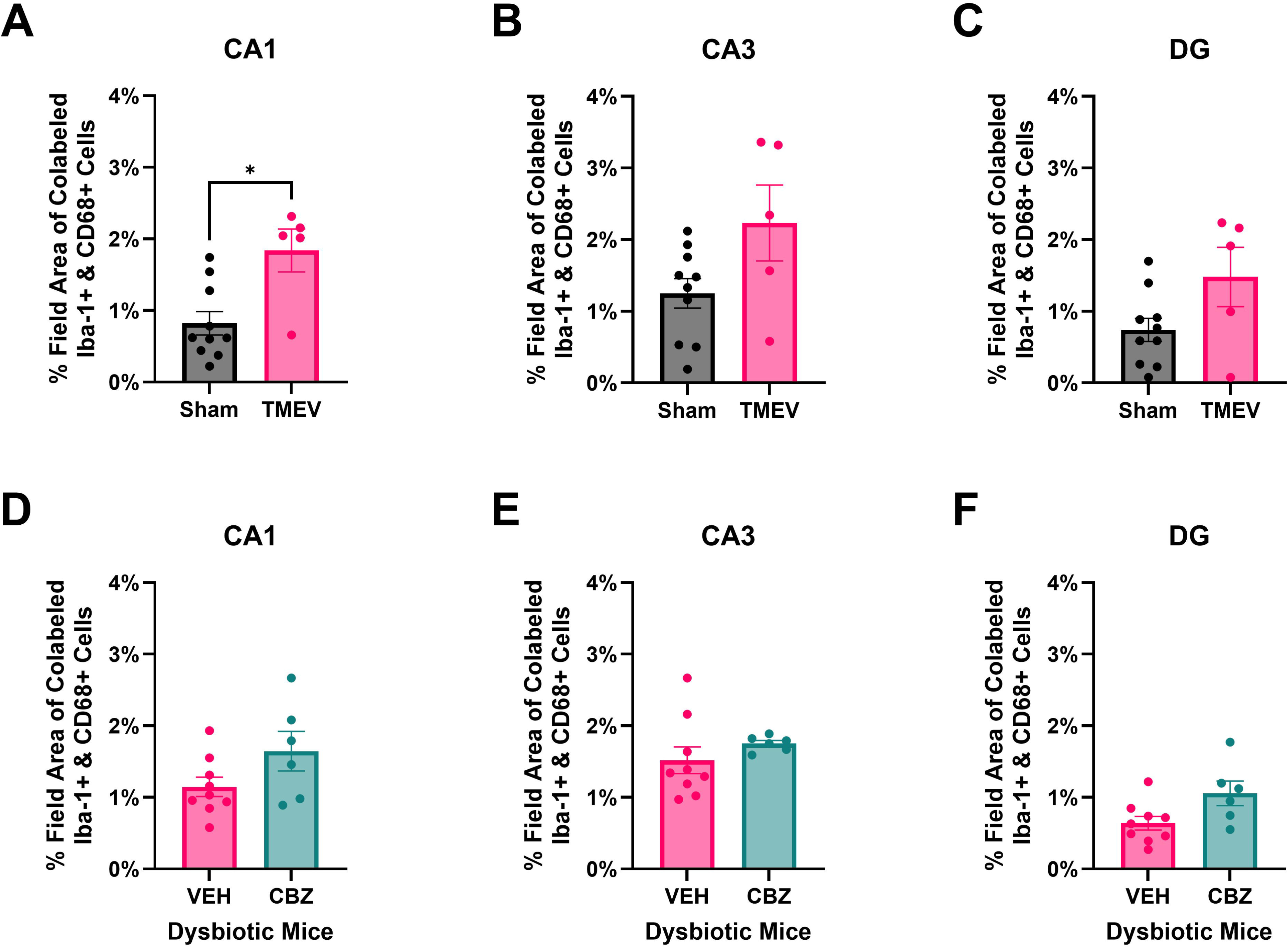
Microglial activation field area changes in hippocampus of male C57BL/6J mice with and without experimentally evoked intestinal dysbiosis (SAL or ABX) 7 days after an acute TMEV or sham infection and subsequent administration of the antiseizure medicine, carbamazepine (CBZ, 20 mg/kg, i.p., bid). All mice, regardless of seizure severity were included in data analysis. A) TMEV infection increased microglial activation in eubiotic mice 7 days post-infection in the CA1 hippocampal subregion. B-C) TMEV infection did not impact microglial activation in eubiotic mice in the CA3 (B) or DG (C) hippocampal subregions 7 days post-infection. D-F) CBZ treatment did not impact microglial activation in the CA1 (D), CA3 (E), or DG (F) hippocampal subregions 7 days post-TMEV infection in mice with experimentally evoked gut dysbiosis. *, *p* < 0.05. Graphs show mean ± SEM.

Astroglial activation, including astrogliosis and proliferation, were assessed across hippocampal regions. TMEV increased total field area with GFAP immunoreactivity, indicative of astrogliosis, in CA1 (F(1,20)=12.14, p<0.01; Figure 5A) and DG (F(1,20)=7.70, p<0.05; Figure 5C), with no significant impact on CA3 (Figure 5B). Conversely, ABX-induced dysbiosis was associated with greater total field area with GFAP immunoreactivity in DG (F(1,20)=5.68, p<0.05). However, CBZ made no impact on astroglia in any hippocampal region. Despite this, astroglia area was reduced in dysbiotic TMEV mice compared to eubiotic TMEV mice in CA3 (F(1,15)=7.24, p<0.05; Figure 5E). *Post hoc* comparisons revealed increased total field area with astroglial marker expression in CA1 in eubiotic TMEV mice compared to eubiotic sham-infected (p<0.05) and dysbiotic sham-infected mice (p<0.05), while eubiotic sham mice displayed reduced astroglia area compared to both microbiome-matched (p<0.05) and dysbiotic TMEV-infected mice (p<0.01). Meanwhile, astroglial proliferation was only impacted in DG whereby CBZ reduced proliferation (F(1,15)=7.08, p<0.05; Figure 6C). Overall, these results indicate that TMEV primarily drives astroglial response in CA1 and DG, while ABX-induced gut dysbiosis differentially impacts this response—increasing it in DG and selectively decreasing it in CA3 in TMEV mice. Notably, astroglial proliferation was primarily affected by CBZ, though this effect was restricted to DG.

**Figure 5.**
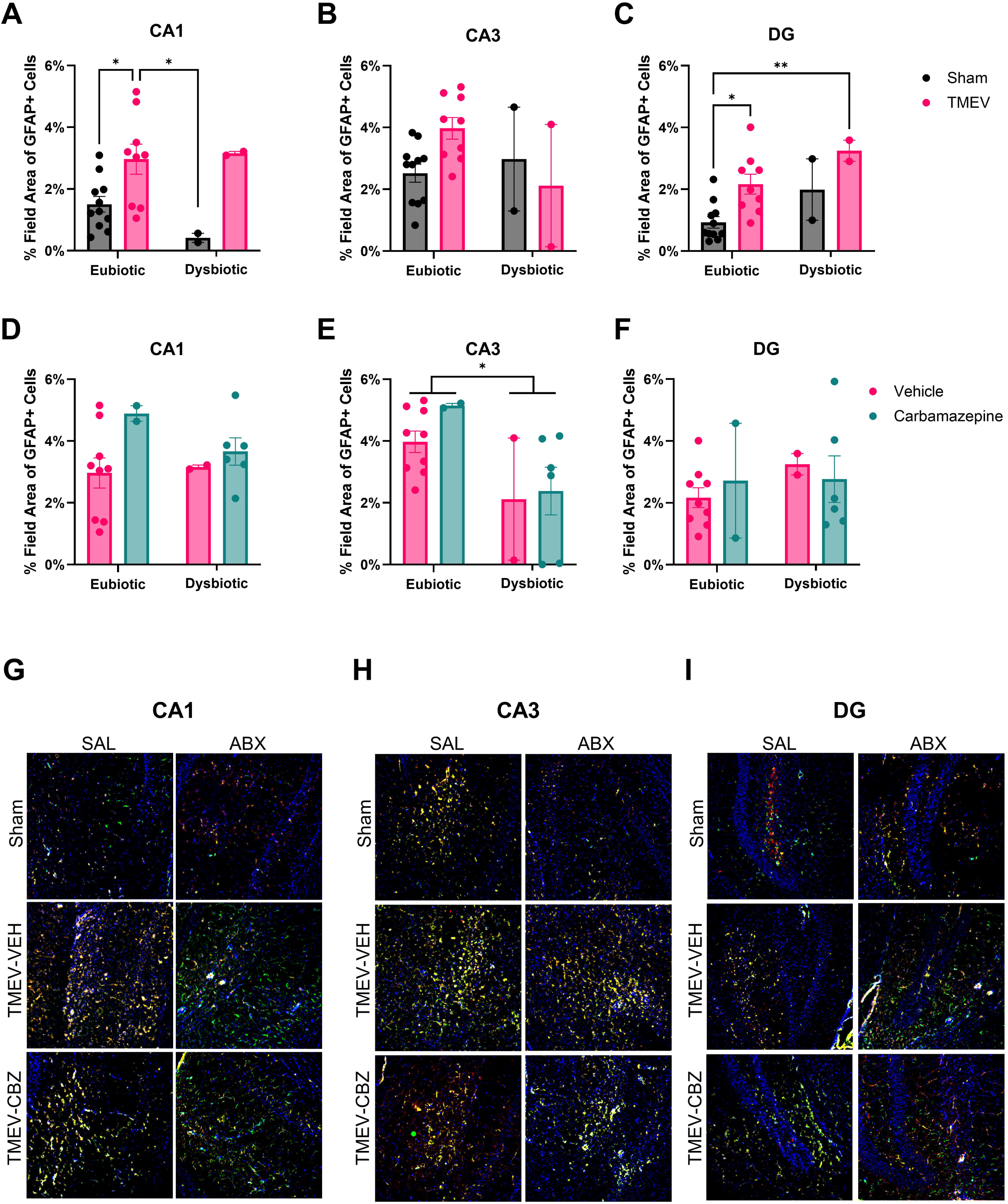
Brain infection with the neurovirulent pathogen TMEV leads to significant changes in the total field area of dorsal hippocampus of male C57BL/6J mice with astroglia marker-positive total field area at 7 days after an acute TMEV or sham infection and subsequent administration of the antiseizure medicine, carbamazepine (CBZ, 20 mg/kg, i.p., bid). Only mice that experienced at least one 3+ Racine seizure following TMEV infection were included in data analysis. A) TMEV infection increased percent field area with astroglial marker-positive signal (GFAP), especially in eubiotic mice, in the CA1 hippocampal subregion 7 days post-infection. B) Neither TMEV infection nor experimentally evoked gut dysbiosis impacted astroglial field area in the CA3 subregion of the hippocampus 7 days post-infection. C) Both TMEV infection and experimentally evoked gut dysbiosis increased astroglial percent field area in the DG hippocampal subregion 7 days post-infection. D) Neither CBZ nor experimentally evoked gut dysbiosis impacted astroglial field area in the CA1 hippocampal subregion 7 days post-infection. E) CBZ treatment did not impact astroglial field area but experimentally evoked gut dysbiosis reduced astroglial field area in the CA3 hippocampal subregion 7 days post-infection. F) Neither CBZ nor experimentally evoked gut dysbiosis impacted astroglial field area in the DG hippocampal subregion 7 days post-infection. G-I) Representative photomicrographs of the CA1 (G), CA3 (H), and DG (I) hippocampal subregions of Ki-67+ (red) and GFAP+ (green) cells with DAPI nuclear counter stain (blue). *, *p* < 0.05, **, *p* < 0.01. Graphs show mean ± SEM, scale bar = 100 µm.

**Figure 6.**
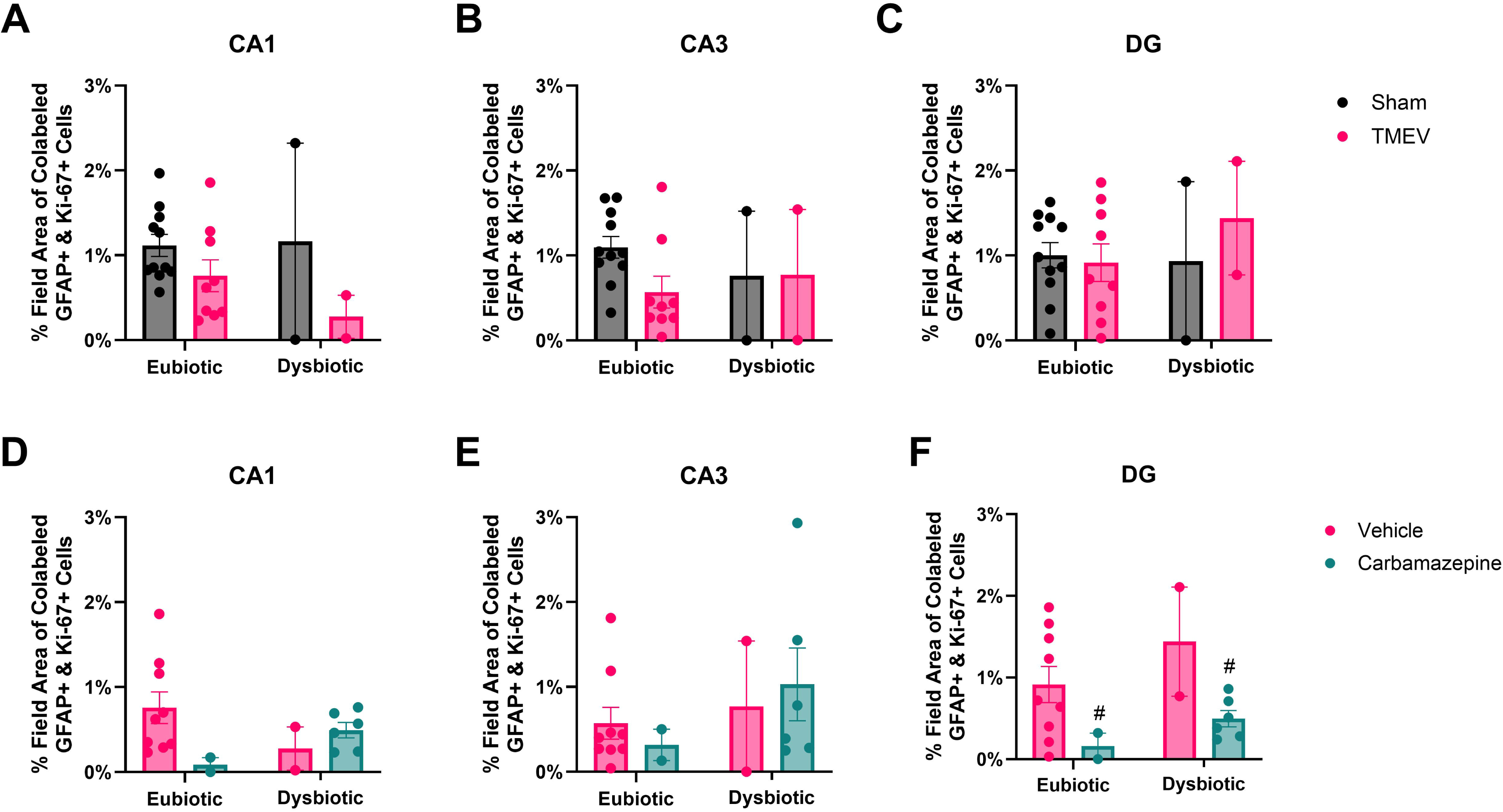
Astroglial proliferation percent field area changes in hippocampus of male C57BL/6J mice with and without experimentally evoked intestinal dysbiosis (SAL or ABX) 7 days after an acute TMEV or sham infection and subsequent administration of the antiseizure medicine, carbamazepine (CBZ, 20 mg/kg, i.p., bid). Only mice that experienced at least one 3+ Racine seizure following TMEV infection were included in data analysis. A-C) Neither TMEV infection nor experimentally evoked gut dysbiosis impacted astroglial proliferation in the CA1 (A), CA3 (B), or DG (C) hippocampal subregions 7 days post-infection. D-E) Neither CBZ nor experimentally evoked gut dysbiosis impacted astroglial proliferation in TMEV-infected mice in the CA1 (D) or CA3 (E) hippocampal subregions 7 days post-infection. F) CBZ reduced astroglial proliferation in TMEV-infected mice in the DG hippocampal subregion 7 days post-infection. #, *p* < 0.05 between CBZ and VEH groups. Graphs show mean ± SEM.

Finally, neurons and neuronal proliferation were assessed across hippocampal regions following TMEV infection, experimentally evoked gut dysbiosis, and the period of repeated CBZ administration. TMEV infection was generally associated with increased neuronal field area in CA3 (F(1,22)=4.54, p<0.05; Figure 7B), while CBZ made no impact in any assessed hippocampal region. *Post hoc* tests revealed that TMEV infection increased NeuN-positive field area only in eubiotic TMEV mice compared to intact sham mice (p<0.05) in CA3. Meanwhile, experimentally evoked gut dysbiosis conferred a presumably greater impact on neuronal proliferation (Figure 8). In CA1, ABX-induced gut dysbiosis was associated with increased neuronal proliferation (F(1,22)=4.54, p<0.05; Figure 8A) and significant dysbiosis × CBZ interactions were observed (F(1,15)=14.42, p<0.01). Conversely, history of TMEV infection was associated with increased neuron proliferation in CA3 (F(1,22)=7.12, p<0.05; Figure 8B) with significant TMEV × dysbiosis interactions (F(1,22)=4.57, p<0.05), but no significant effects from CBZ. DG did not display any significant effects. *Post hoc* comparisons from CA1 revealed increased neuronal proliferation in dysbiotic TMEV mice compared to intact sham mice (p<0.05). When identifying the interaction effects of CBZ and experimentally evoked gut dysbiosis in TMEV mice, ABX-induced gut dysbiosis increased neuron proliferation in VEH-treated mice (p<0.05). This ABX-induced effect was rescued with CBZ treatment (p<0.05). There was a trending increase in neuronal proliferation observed in TMEV mice with ABX-induced dysbiosis in CA3 (p=0.06). Overall, experimentally evoked gut dysbiosis increased neuronal proliferation in TMEV mice in hippocampal CA1, an effect that was rescued by repeated CBZ treatment, while broader impacts on neuronal area were limited to CA3 in eubiotic TMEV mice.

**Figure 7.**
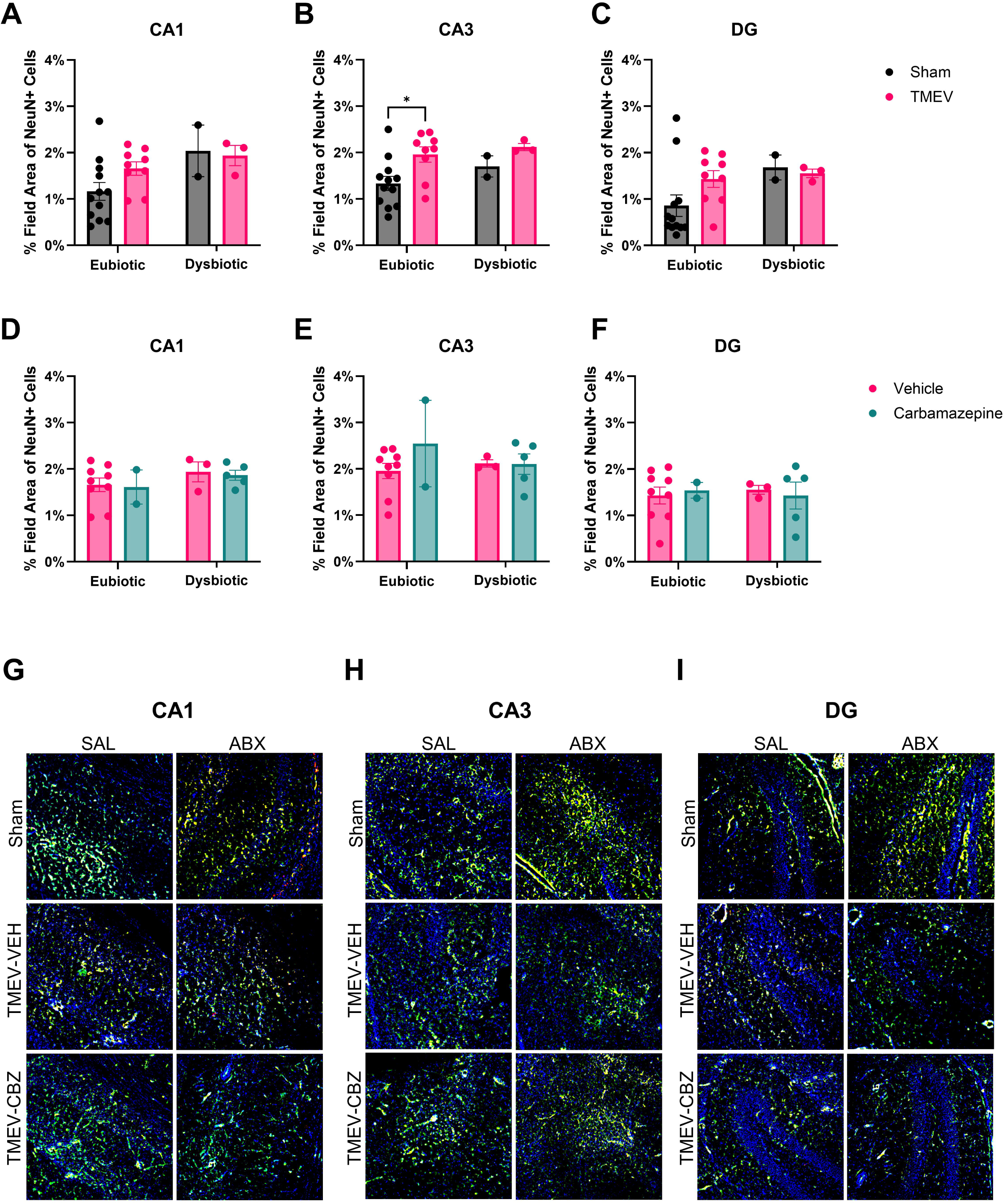
Neuronal percent field area changes in hippocampus of male C57BL/6J mice with and without experimentally evoked intestinal dysbiosis (SAL or ABX) 7 days after an acute TMEV or sham infection and subsequent administration of the antiseizure medicine, carbamazepine (CBZ, 20 mg/kg, i.p., bid). Only mice that experienced at least one 3+ Racine seizure following TMEV infection were included in data analysis. A) Neither TMEV infection nor experimentally evoked gut dysbiosis impacted neuronal field area in the CA1 hippocampal subregion 7 days post-infection. B) TMEV infection increased neuronal field area in the CA3 hippocampal subregion, especially in gut microbiome intact mice, 7 days post infection. C) Neither TMEV infection nor experimentally evoked gut dysbiosis impacted neuronal field area in the DG hippocampal subregion 7 days post-infection. D-F) Neither CBZ nor experimentally evoked gut dysbiosis impacted neuronal area in TMEV-infected mice in the CA1 (D), CA3 (E), or DG (F) hippocampal subregions 7 days post-infection. G-I) Representative photomicrographs of the CA1 (G), CA3 (H), and DG (I) hippocampal subregions of Ki-67+ (red) and NeuN+ (green) cells with DAPI nuclear counter stain (blue). *, *p* < 0.05. Graphs show mean ± SEM, scale bar = 100 µm.

**Figure 8.**
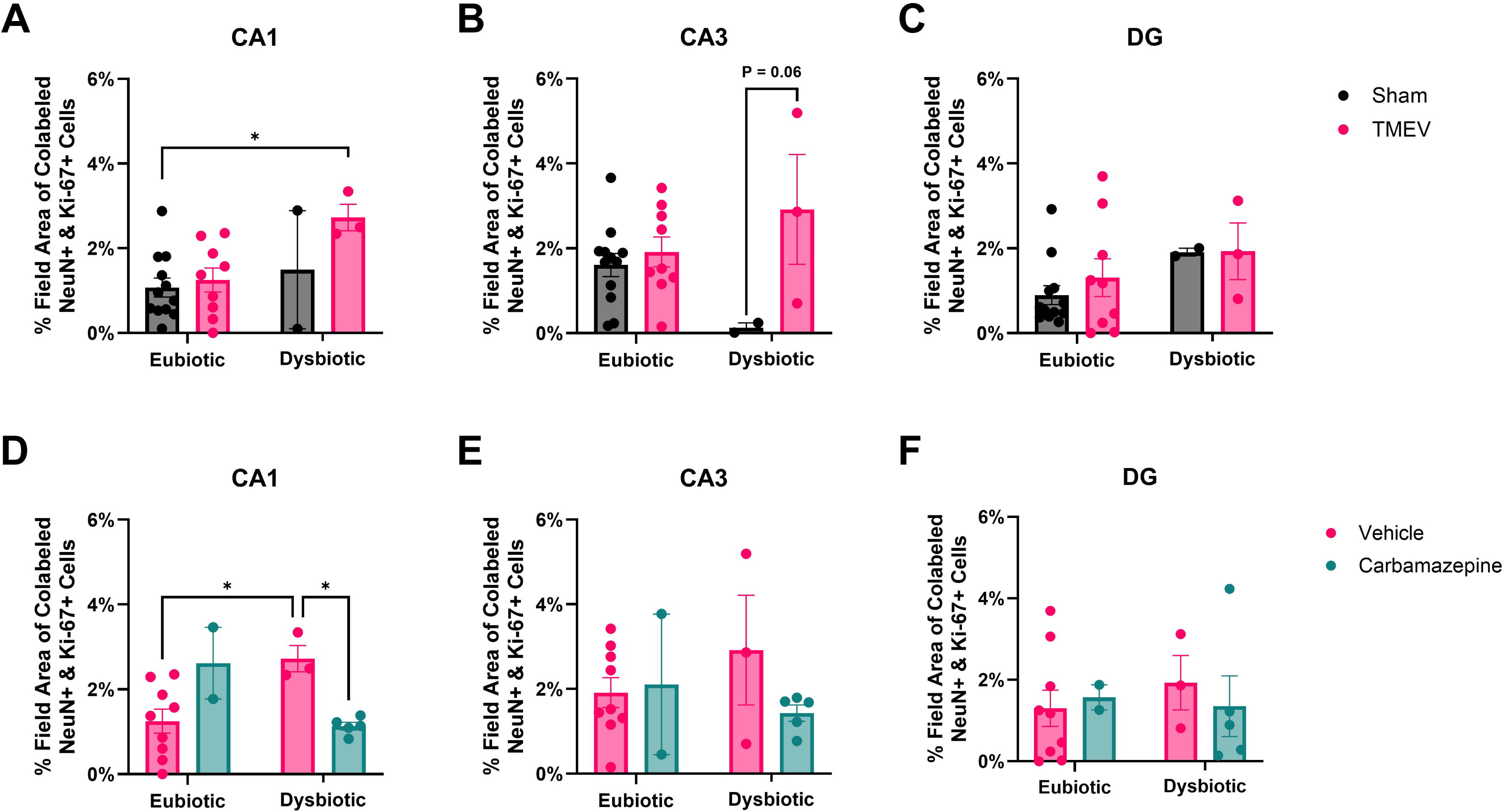
Neuronal proliferation percent field area changes in hippocampus of male C57BL/6J mice with and without experimentally evoked intestinal dysbiosis (SAL or ABX) 7 days after an acute TMEV or sham infection and subsequent administration of the antiseizure medicine, carbamazepine (CBZ, 20 mg/kg, i.p., bid). Only mice that experienced at least one 3+ Racine seizure following TMEV infection were included in data analysis. A) Experimentally evoked gut dysbiosis increased neuronal proliferation in the CA1 hippocampal subregion, especially in TMEV-infected mice, 7 days post-infection. B) TMEV infection increased neuronal proliferation in the CA3 hippocampal subregion, with a trending increase in mice with ABX-induced dysbiosis, 7 days post-infection. C) Neither TMEV infection nor experimentally induced gut dysbiosis impacted neuronal proliferation in the DG hippocampal subregion 7 days post-infection. D) Experimentally evoked gut dysbiosis increased neuronal proliferation in vehicle-treated mice with TMEV but CBZ reduced neuronal proliferation in dysbiotic mice with TMEV in the CA1 hippocampal subregion 7 days post-infection. E-F) Neither CBZ nor experimentally evoked gut dysbiosis impacted neuronal proliferation in the CA3 (E) or DG (F) hippocampal subregions 7 days post-infection.

## Discussion

The TMEV model of infection-induced ASyS is a clinically relevant model of acute seizures and acquired epilepsy that is driven by significant peripheral immune system response, which can itself be shaped by the gut microbiome. This study extends our previous work investigating the behavioral impact of experimentally-evoked gut dysbiosis on ASyS burden and CBZ antiseizure activity shortly after TMEV infection (i.e., 7 days post-TMEV infection; (Erickson et al., 2025)). We now herein confirm our previous findings of neuropathological effects of brain TMEV infection in mice and expand that earlier work to demonstrate that neuropathology is further compounded with experimentally evoked gut dysbiosis coincident with brain TMEV infection. Our present study reveals that experimentally evoked gut dysbiosis leads to exacerbated neuroinflammation (Figure 3). Notably, repeated CBZ administration during the acute seizure period of the TMEV infection did little to counteract this damage, with neuroprotective effects only observed in DG. Importantly, this current investigation provides critical insight into the effects of peripheral immune system priming on CNS pathology in response to a seizure-inducing brain infection, carrying translational potential to managing clinical ASyS and potentially preventing encephalitis-induced epilepsy.

Consistent with earlier findings (Loewen et al., 2016; Umpierre et al., 2014), our results demonstrate that TMEV infection mainly impacts neuroinflammation in hippocampal area CA1 of eubiotic mice. Our results indicate increased microglial and astroglial response, thus also corroborating prior research (Bijalwan et al., 2019; Gerhauser et al., 2019; Stewart et al., 2010b, 2010a). TMEV infection also increased neuronal area and proliferation in CA3 (Figure 7 Figure 8). Neuronal area was specifically increased in eubiotic mice. This result opposes previous research that demonstrates elevated neurodegeneration following TMEV infection (Loewen et al., 2016; Stewart et al., 2010b, 2010a) and opposes our hypothesis that dysbiosis would worsen neuropathology. Moreover, the lack of an effect due to dysbiosis is potentially a result of small sample sizes. Notably, the dysbiotic TMEV group sample size was reduced due to lower incidence of severe seizures (Supplementary Figure 1 and (Erickson et al., 2025)). Thus, the lack of a significant effect may instead be due to reduced seizure severity. Unfortunately there was some unintended tissue loss during the histological procedures that led to the dysbiotic sham group having a smaller histological sample size than was included in our previously reported behavioral study (Erickson et al., 2025). Nonetheless, our present preliminary findings compel future further research to fully elucidate this effect. Meanwhile neuronal proliferation trended towards an increase in mice with ABX-induced gut dysbiosis within CA3, implicating ABX-induced gut dysbiosis as a driving factor. While dysbiosis can impair neuroplasticity (Al Noman et al., 2025; G. Liu et al., 2022; Tang et al., 2021) and contribute to neurodegenerative diseases and psychiatric conditions (Al Noman et al., 2025; Loh et al., 2024; Marano et al., 2023), our small sample size may have limited our results. Thus, dysbiosis-driven effects on neuronal proliferation after TMEV infection warrant further research. Collectively, these findings corroborate previous research showing worsened neuropathological outcomes in CA1 and identifies avenues for further research.

Experimentally evoked dysbiosis primarily affected neuroinflammatory response. First, gut microbiome disruptions led to increased microglial response across hippocampal subregions; microglial proliferation was seen in CA1 and greater field area with microglia signal (Iba-1) was observed in CA3 and DG. Gut dysbiosis alone promotes microgliosis (Çalışkan et al., 2022) and our work further supports this prior report. Other work demonstrates impaired microglial maturation and activation in germ-free, conventionalized germ-free mice, and ABX-treated mice (Erny et al., 2015; Thion et al., 2018). While we did not directly investigate microglial maturation, the elevated microglial proliferation in CA1 suggests that microglial maturation was not impaired under our study conditions. However, astroglial response was hippocampal subregion-dependent. Astroglial response increased in DG of dysbiotic mice, but no effects were observed in CA1. Meanwhile, decreased astroglial response in CA3 occurred in dysbiotic mice, yet this effect only occurred in TMEV infected mice. Dysbiosis itself can cause astroglia dysfunction (Zhao et al., 2021), potentially contributing to the observed inconsistencies across hippocampal regions. However, the astroglial response in DG was further exaggerated by TMEV infection. This is possible due to hippocampal seizures arising from DG, which may act as a brake on neuronal hyperexcitability (Krook-Magnuson et al., 2015; Lu et al., 2016; Petrucci et al., 2025). This is especially notable because we previously found ABX-induced dysbiosis to be anticonvulsant in TMEV mice (Supplementary Figure 1 and (Erickson et al., 2025)). Thus, while mice in this present secondary analysis had a generally lower seizure burden, ABX-induced dysbiosis still contributed to astroglial dysfunction. Overall, these results identify a potential mechanism for the peripheral immune cell migration and activation that drives pathological features in this acute seizure model (Barker-Haliski et al., 2015; DePaula-Silva, 2024; Kirkman et al., 2010; Loewen et al., 2016).

Finally, repeated CBZ administration was investigated to identify its antiseizure impact under dysbiotic conditions and to determine whether this ASM could confer any anti-inflammatory effects in the TMEV model. Prior work reported no effect of CBZ on ASyS severity despite reducing seizure burden (i.e. generalized seizures still occurred, just less frequently), which was associated with no impact on acute astrogliosis, but elevated chronic astrogliosis (Barker-Haliski et al., 2015). Given that diet composition and the associated microbiome modifications on ASyS severity and burden (Zierath et al., 2024), the activity of CBZ was investigated under conditions primed towards inflammation (i.e., experimentally evoked gut dysbiosis). We found mild reductions in microgliosis associated with CBZ administration in TMEV-infected mice, but only in DG (Fig 3F). Within this region, CBZ-treated animals displayed reduced microglial response under dysbiotic conditions, while reduced astroglial proliferation occurred regardless of dysbiosis status. Although prior work reported no impact on acute astrogliosis following CBZ treatment in the TMEV model (Barker-Haliski et al., 2015), CBZ may also possess some anti-inflammatory effects beyond its primary molecular mechanism on sodium channels (Gómez et al., 2014; Matoth et al., 2000; Popović et al., 2025), including direct suppression of microglial inflammatory responses (Boboc et al., 2023; Lauterbach, 2016; Wang et al., 2014). Consistent with this, CBZ mildly attenuated TMEV-induced neuroinflammation in DG under both dysbiotic and eubiotic conditions. These findings are particularly significant given both CBZ’s proconvulsant effects under dysbiotic conditions (Erickson et al., 2025) (Supplementary Figure 1) and the DG’s established role in seizure propagation (Krook-Magnuson et al., 2015; Lu et al., 2016; Petrucci et al., 2025). Thus, despite a higher seizure burden, CBZ retained neuroprotective effects at the site of seizure origin in dysbiotic mice.

Overall, our present study further illustrates that brain TMEV infection in C57BL/6J mice preferentially impacts CA1 over CA3 and DG hippocampal subregions, consistent with earlier reports, and that the intestinal microbiome may play a previously unrecognized role in driving this TMEV infection-induced neuroinflammation in the acute post-infection period (7 days post-infection). However, these findings are limited by the reduced sample sizes, especially in the dysbiotic groups that arose due to technical failures in this secondary study. While the smaller sample size for dysbiotic TMEV mice was due to reduced behavioral seizure severity as reported in our original behavioral study (Erickson et al., 2025), full understanding of the contributions of short-term dysbiosis to long-term epilepsy-related outcomes, as would be possible in TMEV-infected mice, is still needed to better understand how the gut microbiome shapes both acute seizure propensity and long-term epilepsy susceptibility in an at-risk individual. Despite these limitations, our results provide compelling initial evidence that the peripheral immune system may prime brain response in the face of a CNS infection, leading to dramatic shifts in hippocampal network dynamics underlying ASyS-induced neuropathology, which can altogether lead to a difference in long-term seizure-related outcomes. This study demonstrates that the gut microbiome may be a novel contributor to the severity of acute behavioral seizures and associated neurological changes after brain infection, which may warrant GI tract-targeted therapeutic interventions for epilepsy prevention.

## Declarations

### Ethical Publication Statement

We confirm that we have read the Journal’s position on issues involved in ethical publication and affirm that this report is consistent with those guidelines.

### Individual Contributions

Sophia Shonka, Inga Erickson, Melissa Barker-Haliski: conception or design of the work. Sophia Shonka, Inga Erickson, Melissa Barker-Haliski: acquisition, analysis, or interpretation of data. Sophia Shonka, Inga Erickson, Melissa Barker-Haliski: drafting or substantially revising the work.

### Data Availability

All clinical monitoring, behavioral, and histological data is available upon request from the corresponding author.

