## Supplementary Material for "Antibiotic-Induced Gut Dysbiosis Differentially Alters Hippocampal Glial Proliferation and Response After TMEV Infection"

### Supplementary Figures

Supplementary Figure 1. Intestinal dysbiosis with a 10-day course of oral antibiotic cocktail (ABX) disrupts TMEV-induced acute symptomatic seizures (ASyS) presentation from days 3-7 post-infection in male C57BL/6J mice aged 4-5 weeks. Mice were monitored for handling-induced seizures twice per day following administration of the antiseizure medicine, carbamazepine (CBZ, 20 mg/kg, i.p., bid), 30 minutes prior to behavioral seizure assessment. In mice with an intact microbiome, CBZ significantly decreased seizure burden compared to vehicle (VEH). In mice with ABX-induced gut dysbiosis, CBZ significantly increased seizure burden compared to VEH. **, p<0.01. SAL, saline.

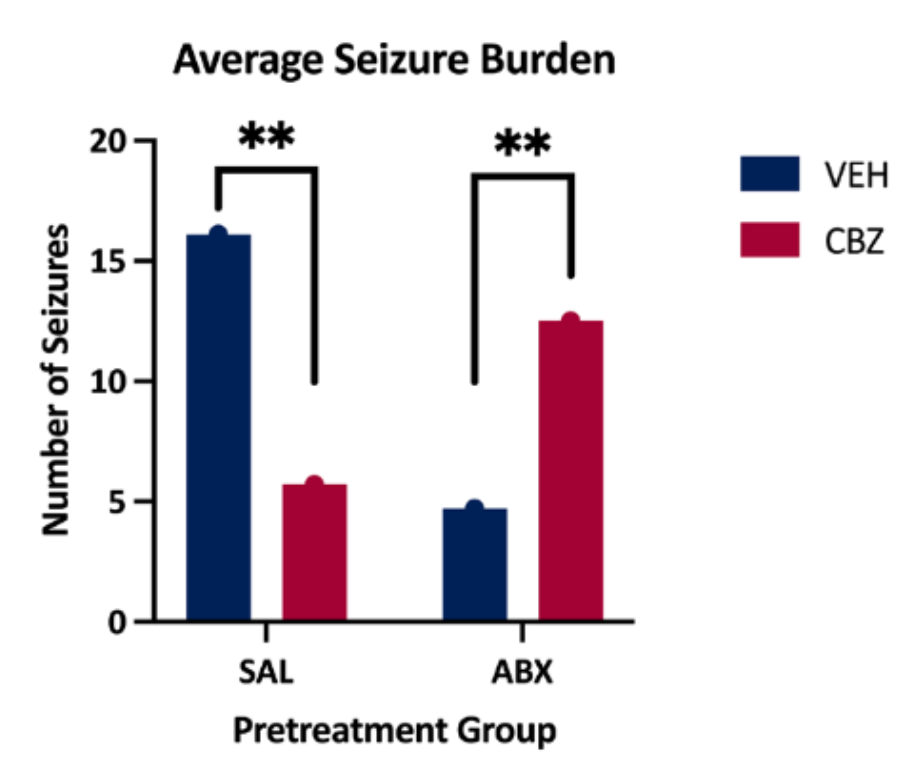

Supplementary Figure 2. A 10-day course of oral antibiotic (ABX)-induced intestinal dysbiosis did not alter plasma carbamazepine (CBZ) concentrations (n=2-5 mice at 0 min; 3-9 mice at 15 min; 3-6 mice at 60 min) in male C57BL/6J mice aged 4-5 weeks at 7 days post-infection with TMEV to induce acute encephalitis and symptomatic seizures. Mice received 20 mg/kg i.p. CBZ twice per day for days 3-7 post-infection, administered 30 min prior to behavioral assessment of handling-induced seizures. On the 7^th^ day post-infection, mice were euthanized within each treatment group for timed collection of terminal blood samples for pharmacokinetic analysis by liquid chromatography/tandem mass spectrometry. Dashed lines represent the anticonvulsant range of CBZ in male CF-1 mice in other seizure tests (Bialer et al., 2004). There were no statistically significant differences in plasma concentrations across treatment groups, as measured by two-factor ANOVA. SAL, saline.

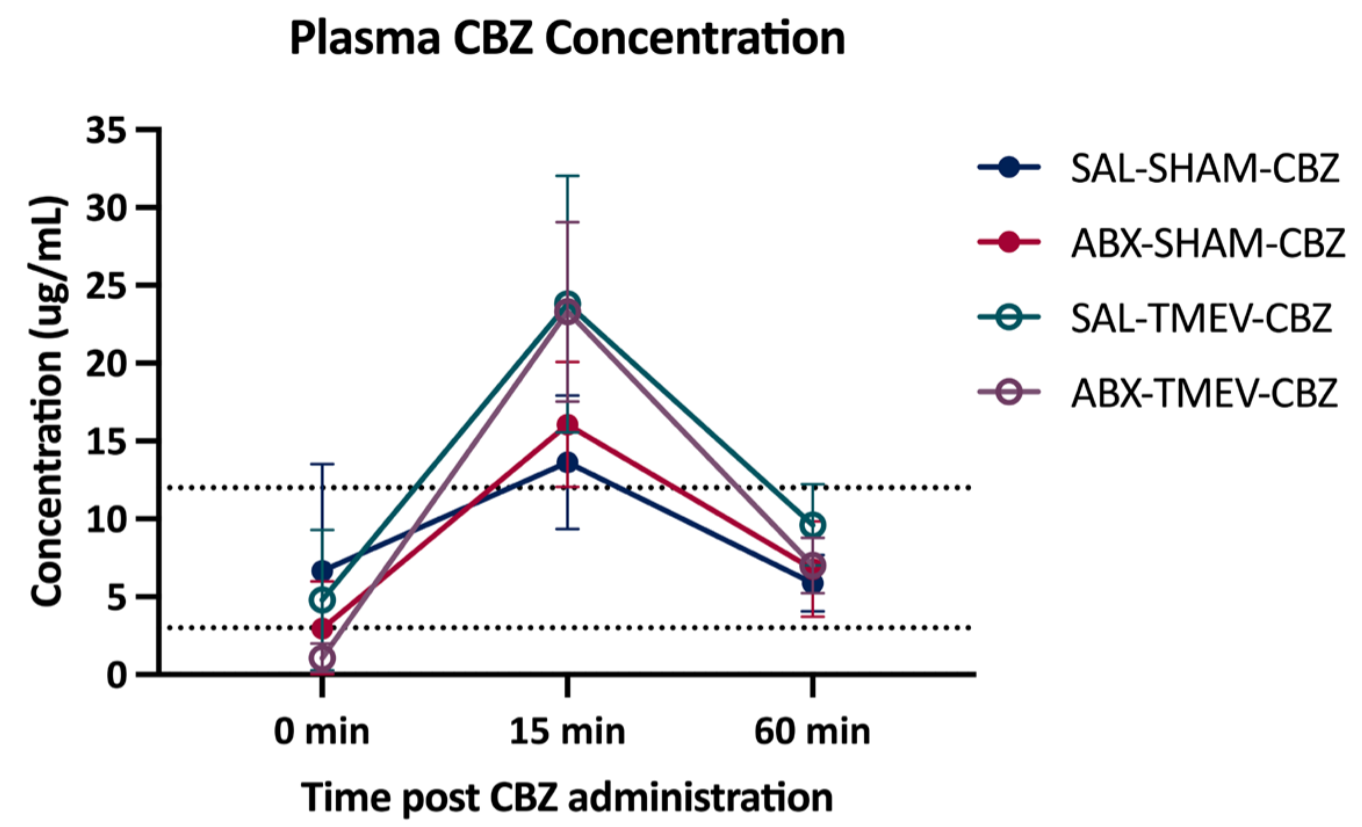
